# LASSIM - a network inference toolbox for genome-wide mechanistic modeling

**DOI:** 10.1101/115477

**Authors:** Rasmus Magnusson, Guido Pio Mariotti, Mattias Köpsén, William Lövfors, Danuta R Gawel, Rebecka Jörnsten, Jörg Linde, Torbjörn Nordling, Elin Nyman, Sylvie Schulze, Colm E Nestor, Huan Zhang, Gunnar Cedersund, Mikael Benson, Andreas Tjärnberg, Mika Gustafsson

## Abstract

Recent technological advancements have made time-resolved, quantitative, multi-omics data available for many model systems, which could be integrated for systems pharmacokinetic use. Here, we present <u>la</u>rge-<u>s</u>cale <u>si</u>mulation <u>m</u>odeling (LASSIM), which is the first general mathematical tool for performing large-scale inference using mechanistically defined ordinary differential equations (ODE) for gene regulatory networks (GRNs). LASSIM integrates structural knowledge about regulatory interactions and non-linear equations with multiple steady states and dynamic response expression datasets. The rationale behind LASSIM is that biological GRNs can be simplified using a limited subset of core genes that are assumed to regulate all other gene transcription events in the network. LASSIM models are built in two steps, where each step can integrate multiple data-types, and the method is implemented as a general-purpose toolbox using the PyGMo Python package to make the most of multicore computers and high performance clusters, and is available at https://gitlab.com/Gustafsson-lab/lassim. As a method, LASSIM first infers a non-linear ODE system of the pre-specified core genes. Second, LASSIM optimizes the parameters that models the regulation of peripheral genes by core-system genes in parallel. We showed the usefulness of this method by applying LASSIM to infer a large-scale nonlinear model of naïve Th2 differentiation, made possible by integrating Th2 specific bindings, time-series and six public and six novel siRNA-mediated knock-down experiments. ChIP-seq showed significant overlap for all tested transcription factors. Next, we performed novel time-series measurements of total T-cells during differentiation towards Th2 and verified that our LASSIM model could monitor those data significantly better than comparable models that used the same Th2 bindings. In summary, the LASSIM toolbox opens the door to a new type of model-based data analysis that combines the strengths of reliable mechanistic models with truly systems-level data. We exemplified the advantage by inferring the first mechanistically motivated genome-wide model of the Th2 transcription regulatory system, which plays an important role in the progression of immune related diseases.

**Author summary:** There are excellent methods to mathematically model time-resolved biological data on a small scale using accurate mechanistic models. Despite the rapidly increasing availability of such data, mechanistic models have not been applied on a genome-wide level due to excessive runtimes and the non-identifiability of model parameters. However, genome-wide, mechanistic models could potentially answer key clinical questions, such as finding the best drug combinations to induce an expression change from a disease to a healthy state.

We present LASSIM, which is a toolbox built to infer parameters within mechanistic models on a genomic scale. This is made possible due to a property shared across biological systems, namely the existence of a subset of master regulators, here denoted the *core* system. The introduction of a core system of genes simplifies the inference into small solvable subproblems, and implies that all main regulatory actions on *peripheral genes* come from a small set of regulator genes. This separation allows substantial parts of computations to be solved in parallel, i.e. permitting the use of a computer cluster, which substantially reduces the time required for the computation to finish.

## Introduction

The ′omics′ era of molecular biology has generated enormous amounts of potentially informative molecular data. However, without new methods for data analysis, this tends to drown researchers in data, rather than provide clear and useful insights. Such methods are considered within systems biology, in particular via its usage of mathematical models [1, 2]. This usage includes mathematical models for predictions of the cellular response to different stimuli, such as a change in environmental conditions or the presence of pathogens. Stimuli are typically recognized by cell surface proteins that induce an intracellular signal transmission via signaling cascades to the cell nuclei. Consequently, transcription factors (TFs) bind to their target genes which results in a change of gene expression [3]. This is reflected by changes in the cellular transcriptome, which may be described mathematically by gene regulatory networks (GRNs) [4]. GRNs consist of nodes representing genes, and edges, representing gene interactions or more general influences[5].

Scientific progress is hindered by the fact that systems biology currently is divided into bioinformatics and mechanistic modeling subfields; the *bioinformatics* approach derives large-scale data-driven networks directly from ′omics′ data, while the *mechanistic modeling* approach performs small-scale non-linear modeling for specific sub-systems. Within bioinformatics, it is common to handle thousands of unknown parameters, aiming at generating a coarse-grained genome-wide view [6-8]. These types of approaches have been shown to perform relatively well in several benchmark studies [9, 10], *e.g.* the LASSO [11-13], ARACNE [14], and the Inferelator [13]. However, these methods do not provide a simulation model that can be used to predict new experiments, and individual model parameters might not be reliable. On the other hand, the *mechanistic modeling* approach typically starts with *a priori* formulated hypotheses regarding a relatively small sub-system representing coupled states. These hypotheses are then formulated into equations, typically non-linear ODEs with unknown parameters that are estimated from experimental data. Hypotheses that disagree with data are rejected, and non-rejected hypotheses are used to predict potentially new regulatory interactions to be tested within new experiments [15-22]. However, mechanistic modeling approaches are greatly limited in size since the simulation-derived parameter estimations scale badly with model size [23]. Because of these inherent limitations in both subfields of systems biology, there are today no methods of realistically predicting system-wide changes in biological systems. For instance, these systems i) are highly non-linear, ii) include critical modeling motifs such as feed-forward and feedback regulations, and iii) eventually lead to genome-wide expression changes. Here we present LArge-Scale SIMulation-based network identification (LASSIM) – a new reverse engineering method of biological systems that resolves i)-iii).

We implemented LASSIM as a toolbox that can handle non-linear ODE modeling of genome-wide processes. The rationale behind LASSIM is that only a subset of all regulators has complex interactions using integrated feedback and feed-forward network motifs with critical internal regulations, while the vast majority of genes are a result of these regulators activity [24]. Moreover, we hypothesize that these regulators can be pre-defined through prior knowledge, i.e. the biological understanding of the system, and these regulators are referred to throughout the article as the *core* system of the specific function. The parameters of the core system can firstly be identified using standard numerical ODE solvers. Once the core system is modeled, the control of peripheral genes can be solved independently. This parallelization of the second part of the network identification enables the run time of LASSIM to linearly scale with the size of the entire genome, which is inspired by the decoupling used in the LASSO method [25]. We have implemented LASSIM as an open source modular Python package using optimized compiled code for computationally heavy algorithmic components. This makes LASSIM fast and flexible enough to handle a great variety of mechanistic and grey-box models.

To show the potential of LASSIM as a flexible and powerful inference tool, describing genome-wide changes from key pathways, we applied it to identify a minimal robust nonlinear model from three kinds of asymmetric data from human Th2 differentiation. This is an ideal model system, since prior work by us and others has shown that the system is transcriptionally controlled by a few relatively well-studied master TFs that orchestrate the expression of a majority of genes [26-31], and that Th2 cells play key roles in regulating the human immune system. We used 12 core TF regulators and, based on DNAase hypersensitivity, we performed binding predictions of the regulatory regions available in Th2 cells, which represented our set of putative interactions. We then applied LASSIM, which performed data fitting and model complexity reduction using time-series gene expression and novel siRNA mediated knock-down experiments during Th2 differentiation such that each TF was directly perturbed once. We assessed the pruned model from randomized subsets of the DNAase predictions in two steps. First, we analyzed the inferred topology of the core system and found that our predicted interactions were significantly better at modeling new Th2 data than other models from the putative interaction list (P<0.006). Second, we analyzed the core-to-gene interactions and found that they were both supported by 14 public ChIP-seq experiments, and identified an enriched set of 685 target genes that had lower cost than 95% of the random models (expected was 385, binomial P<10^−46^). In other words, LASSIM is able to create non-linear minimal models that span the entire genome, and the example presented herein was shown to predict both new data and network structure correctly.

## Results

### LASSIM infers networks from user input

LASSIM is a newly developed method for large-scale flexible non-linear ODE modeling, with an accompanying implementation built as a Python open source software package (presented in Fig. 1), and can be found at https://gitlab.com/Gustafsson-lab/lassim. The rationale behind LASSIM is the empirical observation that many biological processes are controlled by few (less than 50) interlinked regulators that includes feedbacks, hereafter referred to as the *core system.* LASSIM is built to first solving the core system and thereafter solving each other peripheral gene in parallel. This significantly reduces the running time (Fig. S1). LASSIM has been implemented modularly, allowing for methods and algorithms to be easily exchanged, and is built on PyGMO, which is the European Space Agency platform for performing parallel computations of optimization tasks [32].

**Fig 1.**
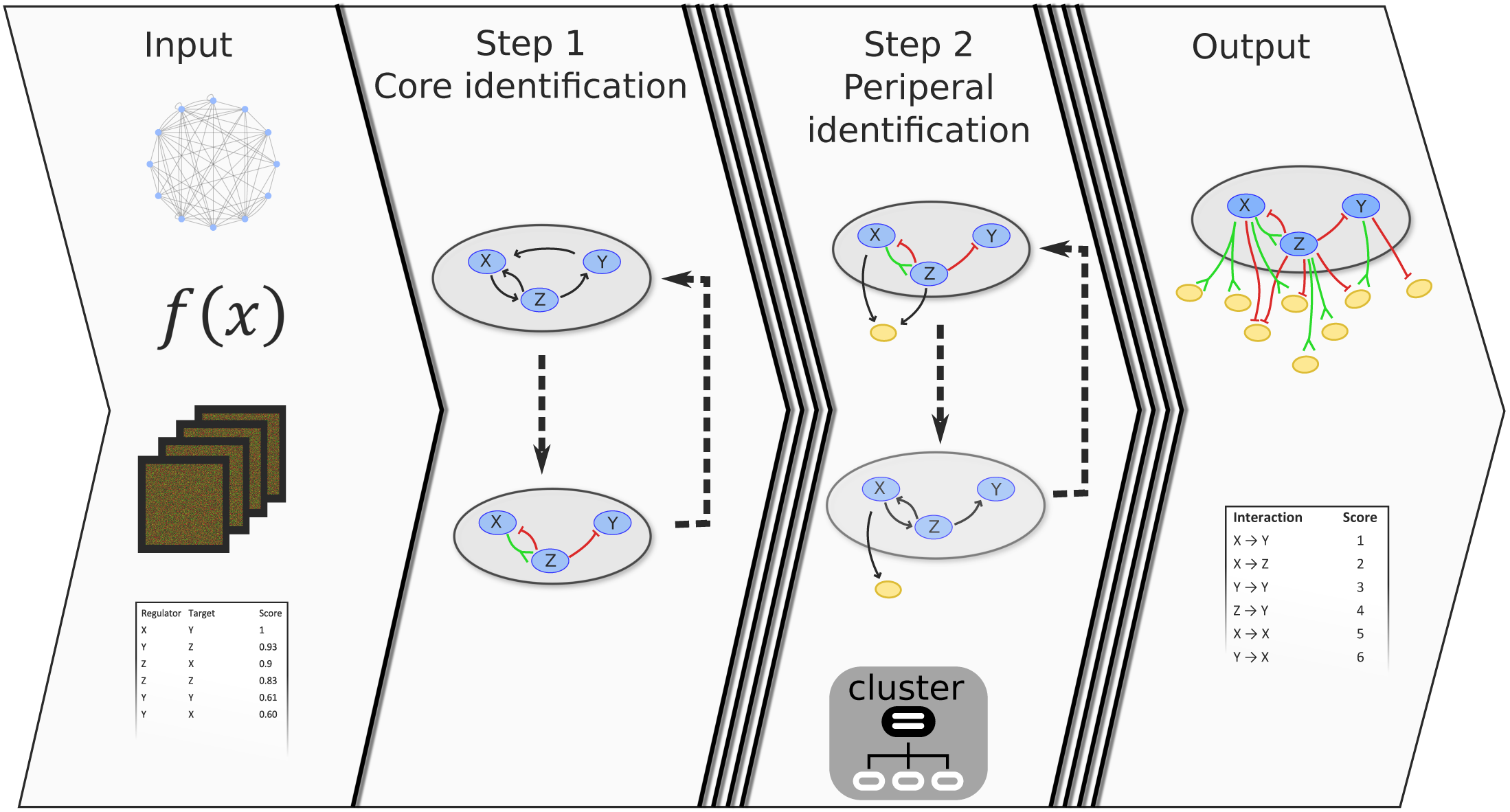
General workflow of the LArge-Scale SIMulation (LASSIM) high performance toolbox. LASSIM requires four basic inputs (left), i.e. a core and peripheral network, expression data and dynamic equations, and fits a large-scale non-linear dynamic system based on the fully parallel, modular, ultra-rapid global optimization toolbox PyGMO developed by ESA. First, LASSIM performs pruning and fitting of a core dynamic system, and then expands it by adding peripheral genes in parallel using a computer cluster. The LASSIM functions are fully modular, and have been built so that the functions describing the optimization procedure, dynamic equations, cost function and data pruning can easily be changed by the user.

LASSIM takes as user input the *a priori* structure of a system, dynamic equations, model selection criteria, expression training data (time-series and/or perturbation data), and a confidence interaction list (Fig. 1, Input). The default kinetic equations are built on a sigmoid activation component and a linear degradation component according to the results presented in [33, 34]. However, linear, Hill, mass-action, or other user defined kinetics can easily be incorporated thanks to the design of the LASSIM interface, by simply replacing default computational components with kinetics coded as Python functions. The models are by default tuned via backward selection and model selection by minimization of likelihood ratio-tests. LASSIM starts by inferring parameter values for the internal regulations within the core system (Fig. 1, Step 1), and unnecessary parameters in the core system are removed using training data such as response to perturbations and time-series measurements. In a second step, the core-to-peripheral gene regulations are modeled independently, taking advantage of the parallel nature of the problem, again with model selection based on a user-defined criterion (Fig. 1, Step 2). Finally, due to their independence, the combined models are assembled into one single full-scale non-linear ODE model (Fig. 1, Output). We used two data generated examples to illustrate that including realistic dynamics provided both improved network structure identification and trajectory modeling compared to a pure static regression-based LASSO approach (Fig. S2).

### LASSIM modeled Th2 differentiation of naïve T-cells

We next applied LASSIM to infer a genome-wide mechanistically motivated model of TF-target relationships of human differentiation of naïve CD4+ T-cells becoming effector Th2 cells, which play a pivotal role in multiple immune-related processes (Fig. 2A) [35]. We first identified a putative core set of 12 TFs (COPEB, ETS1, GATA3, IRF4, JUN, MAF, MYB, NFATC3, NFKB1, RELA, STAT3, USF2) from our previous experimental studies of TFs within the context of Th2 differentiation [35, 36], and a set of 11,083 differentially expressed genes in fully developed Th2 cells vs naïve T-cells (FDR<0.05, see Methods). Second, we predicted 63 core TF-TF putative interactions and 64,872 putative peripheral interactions with Th2-specific DNase-seq footprints using the HINT bias correction method (Fig. 2B) [37, 38], together with motif matching from three TF binding motif databases UniProbe, JASPAR, HOCOMOCO using the regulatory genomics toolbox Python package rgt-gen [39-41]. We then trained LASSIM on our previous microarray time-series data (Fig. 2C) from naïve T-cells to differentiated Th2 cells [35], and a compendium of siRNA mediated knock-downs under Th2 differentiation (within 16-24 h) of each of the respective TFs analyzed with microarrays (Fig. 2D). We compiled the compendium re-using six previously published siRNAs and created six new siRNAs (Methods) together with 18 data points (each series repeated 3-4 times), for which we applied LASSIM. The initial core system comprised of 216 data points and 75 parameters, for which we applied LASSIM to perform a fitting of the core system to the time-series data and all siRNA knock-down data simultaneously (Fig. 2C-D). The corresponding starting peripheral systems had 18 data points and about eight parameters per gene.

**Fig 2.**
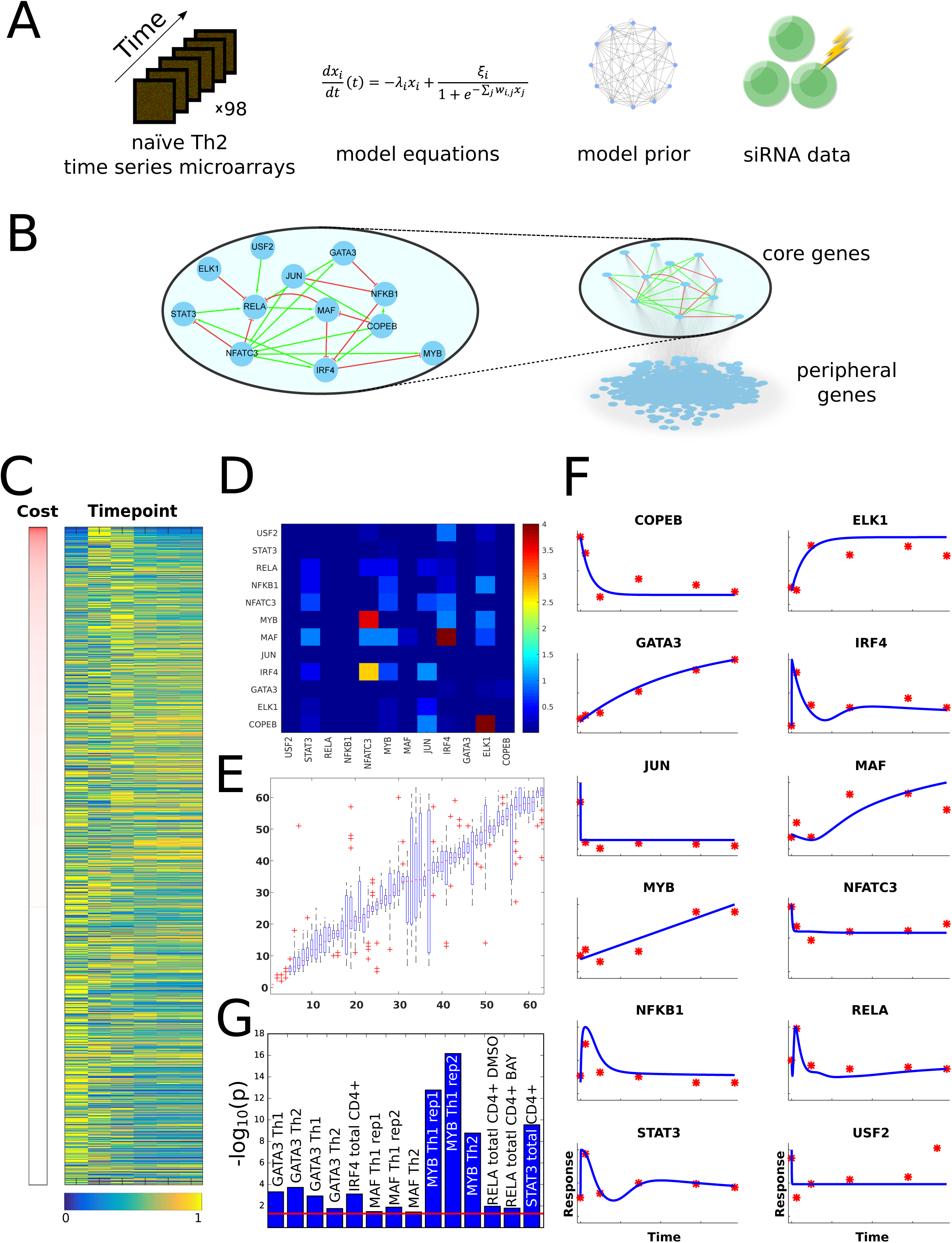
LASSIM inferred a robust minimal full-scale non-linear transcription factor - target dynamic system describing naïve T-cells towards Th2 cells. (A) We identified 12 core Th2 driving TFs from the literature, inferred their putative targets using DNAase-seq data from ENCODE. This was, together with Th2 dynamics, and siRNA of each TF followed by microarrays, used by LASSIM to infer a Th2 system, which is shown in (B). (C) The measured mRNA profiles of the 10 543 peripheral genes (blue/yellow represent relative low/high expression), sorted by the model cost (*V*(***p****\**)). (D) Heat map of the data fit of the Th2 model to the siRNA perturbation data, i.e. the siRNA part of *V*(***p****\**). Each siRNA knock experiment is represented as a separate column. For example, the model fits the response of a knock-down on RELA well, but fails to fit the response of MAF when IRF4 is knocked down by siRNA. (E) Box-plot representing the ranking of each removed parameter from multiple runs of LASSIM from somewhat different starting positions. All edges that had a median over 39 were included in the final model. (F) Microarray experiments (red dots) and simulated dynamics of the model (blue solid lines) of the core TFs. (G) The results from the ChIP-seq analysis, where the y-axis shows the-log_10_(P). The red line denotes the significance level 0.05.

### Robustness analysis identified a LASSIM model with a minimal number of parameters and low error

To estimate the robustness of our fitted model we repeated the full LASSIM procedure, including stepwise removal of edges in the core system 100 times using different starting points and small perturbations of the data (Fig. 2E, S3-4). We found robust minimal solutions at 40 removed parameters, simultaneously adhering to the following characteristics: 1) Good fit to the time-series data, i.e. non-rejected solution from a Gaussian noise distribution (Eq. 3 in Methods gives cost function *V*(*p\**) = 15.0, χ^2^_0.95_ (df = 12*6) = 92.8) (Fig. 2F). 2) Good fit to the siRNA data (Fig. 2D). 3) Robust sets of interactions, i.e. when LASSIM was applied to the same training data starting from different initial parameter sets the orders of parameter removal were highly similar (Fig. 2E).

The problem of model selection and parameter estimation is a complex optimization one due to the large number of variables and the non-convex nature of the objective function. We therefore analyzed the sensitivity of the core system, and found the error to increase more through bootstrap re-sampling than between re-runs of LASSIM on the original data (Fig. S3), and that removal of all siRNA data increased the model variance (Fig. S4). These two results supported the system with 40 removed parameters, i.e. the 23 remaining interactions, as a robust minimal core model.

### LASSIM identified peripheral interactions that were functionally supported by ChIP-seq

Next, LASSIM stepwise pruned the interactions and fitted the parameters representing the core regulation onto the peripheral genes. There was a good agreement between model simulations and time-series measurements (i.e. *V*(*p\**) < χ^2^_0.95_ (df = 6) = 12.6) for 10,546 genes (Fig. 2C), as well as for the 12 siRNA knock-downs (Fig. 2D). The resulting genome-wide model consisted of 35 900 core-to-peripheral gene interactions, thus removing 28 972 potential interactions. In order to evaluate the performance of LASSIM we tested whether the removed or remaining interactions were more likely to be physical interactions. This was done for each individual TF by comparing the mean peak scores of the removed and remaining interactions respectively using ChIP-seq and ChIP-Chip from 14 experiments of five different TFs. [35, 42-44]. We found consistent enrichments for the remaining interactions (permutation test P<0.05) for each experiment, with the highest enrichments from the STAT3 and MYB bindings (P<10^−9^, see Fig. 2G). Thus we feel confident that the minimal model robustly removes interactions that are less likely to be direct than the remaining interactions.

### LASSIM monitored total CD4+ T-cell dynamics significantly better than other models from the prior network

In order to test the general applicability of the inferred Th2 dynamic model we aimed to test its capability to model Th2 differentiation from a slightly different starting point. Th2 cells were therefore differentiated *in vitro* starting with a mix of naïve, memory and primary CD4+ T-cells, hereafter called the *total* T-cell population, which through their respective cytokine secretion influenced the process. We first asked whether the model developed for Th2 differentiation of naïve CD4+ T-cells could also monitor that of total T-cells. For this purpose, we performed new time-series experiments of four time-points and 15 microarrays (Methods). We reasoned that as the new cell-type contained also memory and primary cells the magnitude of the kinetic parameters might change, but we hypothesized that the underlying interactions and signs should not change. We therefore retrained the identified model as well as randomly sampled models from the TF binding prior network and of the same size and parameter sign distribution as the minimal Th2 model. The retraining was performed similarly for all models using the optimization procedure of LASSIM, and our test statistic was the cost functions *V*(*p*). We found a good re-fit for the core system, which passed a χ^2^-test *V*(*p\**) < χ^2^_0.95_(df = 4*12) = 65.2) (Fig. 3B) and 9 992 peripheral genes (Fig. 3C). Then, we compared the refitted core model against 1000 null models, which showed that its residual was lower than all except six null models (bootstrap P = 0.006, Fig. 3D). To analyze whether these random models represented missed good solutions by LASSIM, we retrained the kinetic parameters of these six models on our first training data, but could not find any random models with as low cost as the identified minimal Th2 model (Fig. S5).

**Fig 3.**
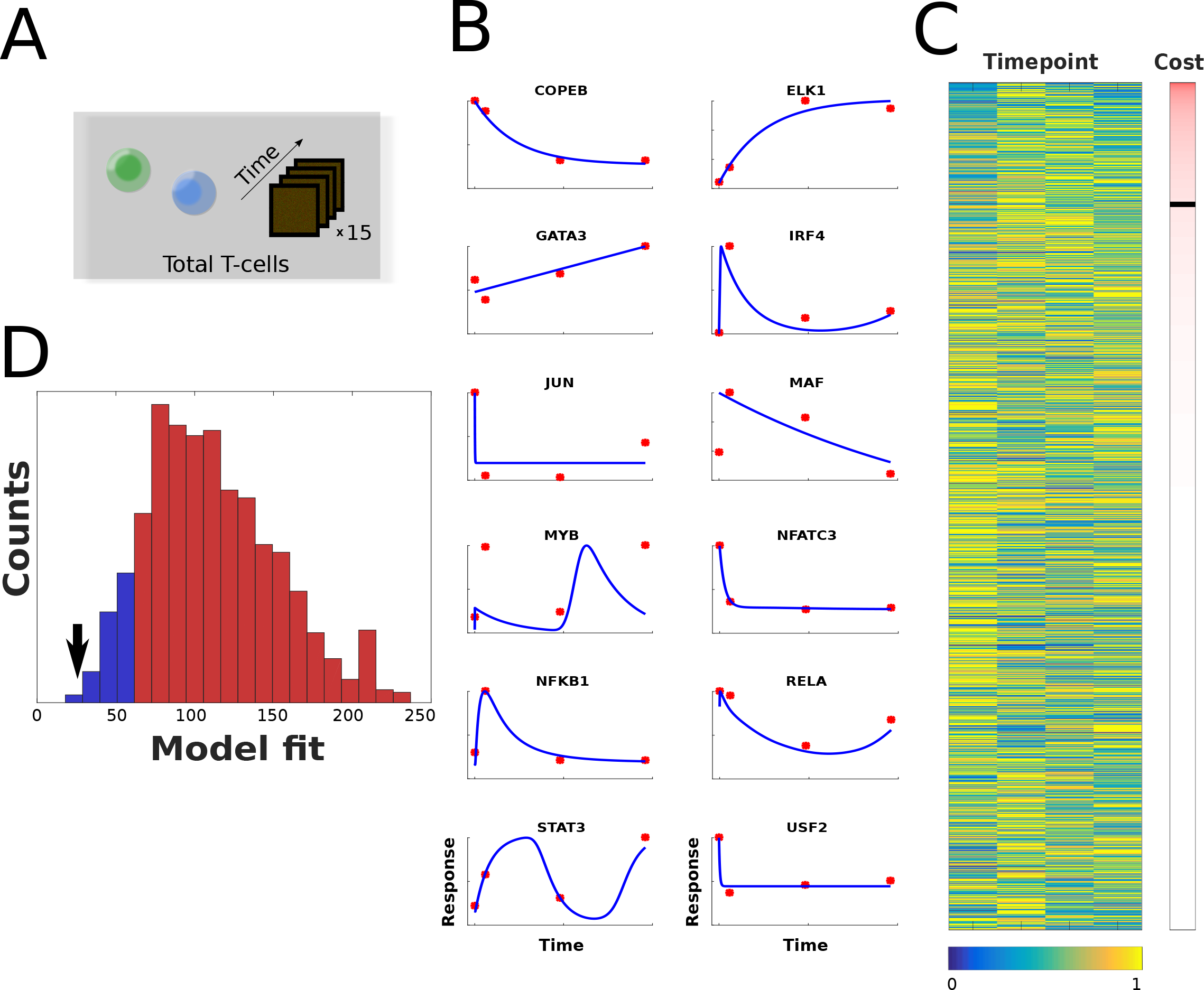
The edges from the naïve Th2 model were better refitted to total Th2 differentiation than the prior network. (A) We refitted the core network to new time-series data from total Th2 by keeping the signs of the inferred minimal Th2 model. (B) The fit of the core model to the total T-cell model is shown. The y-axis has arbitrary units, and the x-axis is time. The model output is the blue curves, and the data are the red points. (C) The fit of the peripheral genes. The figure follows the same style as in Fig. 2F. The black band in the cost bar denotes the 0.95 rejection limit of the corresponding χ^2^-test, with all rejected models above the band. (D) We resampled our prior matrix with the same number of parameters and signs as the Th2 model 10,000 times, and compared the random models with the output from LASSIM. The ability of the inferred core network structure to fit novel gene expression data of total T-activation was considered. A total of 1 000 random core model structures were drawn from the prior network, and refitted to the data. The distribution of the fit is shown by the blue curve. The core network identified from LASSIM was showed to be significantly better for fitting the novel data, as marked by the black arrow. Moreover, most of the core models could be rejected by a χ^2^-test, and are represented in red.

Lastly, we proceeded to a similar test for each of our peripheral genes by identifying 100 random models for each gene (i.e. one million random models, Fig. S6). Many peripheral genes had high model complexity both in the random distribution (average in-degree = 5.9) and in the minimal model (average in-degree = 3.4), thus we had low statistical power and could not perform stringent statistical corrections for testing which of the peripheral genes were significantly better than random. However, for 64% of the peripheral genes our model performed better than the average random model (binomial test P<10^−130^), and for 685 genes our model performed better than 95% of the corresponding random models (expected 385 genes, P < 10^−46^). In summary, we conclude that the LASSIM identified core and peripheral solutions could monitor new dynamic data relatively well.

## Discussion

Gene regulation constitutes a complex dynamic system that can drive cells to new cell states during differentiation. These systems are often analyzed using mathematical modeling via *bioinformatics* and *mechanistic modeling,* respectively. Although both approaches have distinct advantages, few efficient ways of combining them have been identified. Herein, we have presented the LASSIM toolbox, which is built on a small-scale simulation-based framework, and it is therefore theoretically possible to model any combination of standard biochemistry kinetics, including classic mass-action and synergistic Hill-kinetics. LASSIM is built on the assumption that a small-scale *core* model can be found–which is regularly assumed with the small-scale modeling community. The core model is applied as input to the rest of the *peripheral* genes, and this process is computed in parallel, under the assumption that the there are no feedback interactions from the peripheral genes to the core genes. The outcome of LASSIM is a full-scale dynamic gene regulatory model.

Although LASSIM is different to other GRN inference methods, it is useful to compare close alternatives. ARACNe is a frequently used fast information theoretical method that produces genome-wide networks [14]. However, it cannot be used to predict the directionality or sign of interaction, and cannot be used to predict the dynamic response upon treatment due to the lack of model equations. Model equations are present in few ODE-based models, such as Inferelator [13], ExTILAR [45], and NetGenerator [46]. Inferelator and ExTILAR are directly based on regression, thus they use estimated time derivatives directly from data to approximate ODEs. ExTILAR [47] use target genes to infer upstream TF activity and solve a fully coupled regression problem of TF-targets explicitly. After identifying the structure of a model by regression, ExTILAR fits ODEs to that structure to achieve a model, which can be simulated. However, the tool can only predict models of moderate size but not full genomic systems. Inferelator has been built to include great variety of user-inputs, and some different kinetics, while ExTILAR uses linear models. However, neither of the methods can handle mechanistic models with feedbacks. There are several tools for small-scale mechanistic models, where NetGenerator [46] represents one of those commonly used for GRNs. This method is based on linear or non-linear ODEs, and makes use of simulation-based models to estimate parameters. NetGenerator uses a heuristic search strategy with forward selection and backward elimination to infer a sparse network and can, similarly to LASSIM, use several perturbations simultaneously for parameter estimations and model selection. However, this method is not optimized for large-scale systems, and practical examples are limited to some tens of genes. In conclusion, there is to date no other approach that allows for the identification of a GRN simulation model for the entire genome.

The key assumption behind LASSIM is that all genes are affected only by a core system. For the specific example considered here, Th2 differentiation, the studied core genes have already been proposed and studied in previous works [27, 28, 35, 48]. More generally, it is reasonable to assume that the most influential TFs can often be considered as a sub-system, which can qualify as a core-system in the LASSIM sense. The two main arguments for this are 1) that as the number of components, i.e. genes, required to sustain a biological function, grows, the easier it becomes to destroy the function [24], and 2) that the interactions within the TF core system can be considered as collapsed versions of more extensive pathways. 1) can be understood by considering that each component works with probability p, then the probability that the function will be performed decreases with the number of components n as p^n^. For instance, if p=0.99 and n=10, then the function is performed in 90% of the cases, but if n=1000, then it is only performed in 0.004% of the cases. A core system with relatively few crucial components should therefore be behind each function. 2) can be understood by considering two hypothetical TFs in a core system: **TF**A and **TF**B. Assume that **TF**A affects **TF**B. but via one other gene C outside the core-system. In other words, the true pathway is **TF**A −> C −> **TF**B. In LASSIM, this interaction would still be captured, but as **TF**A −> **TF**B. i.e. as a collapsed interaction. Such collapsed interactions form the basis of GRNs as such, since GRNs describe the collapsed effect of a gene directly on another gene, even though this effect may in reality be indirect, occurring via, for instance, phosphorylation of proteins. The existence of a core system is therefore likely to be an effective simplification. Note that, in general, a different core system is behind different biological functions, so it is necessary to take special care when combining data from different biological functions.

The application of LASSIM on Th2 differentiation presented herein is of principal importance since it is the first simulation-based model developed for the entire genome, and integrates three different kinds of data, i.e. prior network estimates, time-series, and knock-down data to infer a robust minimal model with realistic dynamics. The first principal requirement was a core system, where we relied on our previous findings such that a majority of core genes originated from Bruhn et al [36], which is a practical example of how a core system could be *a priori* identified. They identified a Th2 module and its upstream TF regulators, where we consideed the regulators as core genes. We applied LASSIM to infer a simulation based model of this system, based on Th2-specific binding predictions. Our analysis showed that about 45% of the TF-target predictions could be removed from the model with a similar fit to data. Importantly, the removed interactions were also the ones with the lowest experimental support from ChIP analyses of the individual TFs, which shows support for the inferred peripheral gene regulations. We also tested whether the model was capable of modeling new expression dynamics from mixed cell differentiation. For that purpose, we had to adjust the magnitude of the kinetic parameters. We therefore compared the LASSIM-inferred model against the random models sampled from the TF-target prior predictions and allowed for a similar retraining as for the Th2 model. We found both that the core model and an enriched fraction of the peripheral genes were significantly better modeled by LASSIM. Interestingly, the core TFs MAF and MYB yielded the worst fit to the dynamic curves, but still significantly overlapped with ChIP-seq (**P**_MAF_ <0.04, ._MYB_ <2*10^−9^). These observations suggest that our modeled dynamics might reflect activation from post-translational modifications of MAF and MYB, where, for example, tyrosine phosphorylation of c-MAF has been shown to increase IL4 and thereby amplify Th2 differentiation in mice [49]. From an experimental design point of view, the LASSIM output also demonstrated the need for perturbation data of all regulators in the studied core system in order to have enough information in data for making model claims. Moreover, by analyzing the underlying motifs in the core model, we can observe feed-forward and feedback loops, e.g. the feedback STAT3-RELA-MAF-IRF4-STAT3 path. Hence, LASSIM ODE models are needed to perform pharmacokinetic modeling of drugs. In summary, LASSIM enables the large-scale modeling of several data types including knock-downs and phosphorylation time-series, for which mechanistic non-linear models are needed, for example to include mass-preserving constraints.

## Methods

### Selection of core system TFs and predicted interactions

Several TFs have been tested for being associated with Th2 differentiation. We base our selection on the following four studies: 1) In Bruhn et al. [36], we performed siRNA of 25 TFs and found seven highly relevant (GATA3, MAF NFATC3, STAT5A, STAT3, NFKB1, JUN). 2) In Gustafsson et al. [35] we also identified the TF MYB as having great importance to Th2. 3) We have also identified IRF4 and ETS1 as key TFs of Th2 differentiation [43]. 4) By analyzing data from Nestor ***et al*** [50] we identified USF2, and KLF6 as potentially interesting Th2 TFs because they were differentially expressed in unstimulated non-symptomatic allergic patients vs controls (P_USF2_ = 3 * 10^−9^, P_KLF6_ = 0.001).

### Model equations

Systems of ordinary differential equations (ODEs) are frequently employed in engineering to mathematically model the states of a system. Let the *i*:th state be denoted *x_i_*, and let all states be collected in a vector ***x***. The dynamics of the states are given by ODEs (Eq. 1), which describe the changes, i.e. using a function ***f*** of states ***x*** and additional parameters ***p***.

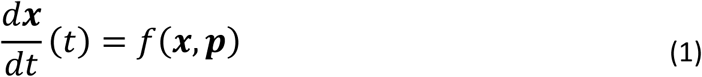

Simulation-based approaches, like LASSIM, can be developed for any form of f, e.g. nonlinear Michaelis-Menten or mass-action expressions. For gene regulation mechanisms, however, a more realistic form of ***f*** is a sigmoid regulation, used in e.g. [33, 34], as shown in Eq. (2):

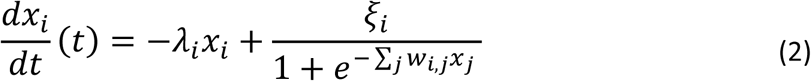

In Eq. (2), the degradation rate of gene *x_i_* is negatively dependent on the concentration, while the regulators *x_j_* can have both inhibitory and enhancing effects. The rate constants are *λ_i_, ξ_i_*, which are strictly positive, and *w_ij_*, which can be both negative and positive. Given this equation, the value of the second term is restricted between 0 and *ξ_i_* and the equation system is stable for all *λ_i_, ξ_i_∊* ]0, ∞[. The search bounds of the parameters were defined to be *λ_i_, ξ_i_∊* [0,20], *w_ij_ ∊* [-20,20] which covers almost all of the possible values of the term (Fig. S7).

### Model selection

When fitting the model to data, the objective function was set to minimize the sum of squares.

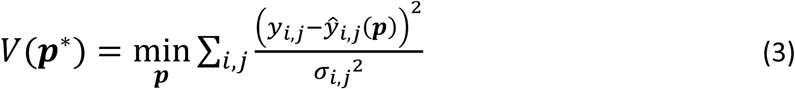

In Eq. (3), the best parameter vector, ***p****\*,* is the parameter set that minimizes the difference of the measurements y and model output *ŷ* for state *i* and time point *j.* For the knock-down data, the fit was calculated correspondingly, but with the standard deviation σ set to 1. This selection of σ made the cost contributions *V*(***p***) from the knock down and time simulations to be of the same order of magnitude for most parameter sets. For the time series data, standard deviations were estimated using all time points. Both the perturbation and time series model output were fitted using Eq. (3). Moreover, the structure of Eq. (3) can be used to fit any type of model output, e.g. dose-response curves, knock-in, or steady state data.

The model selection process in LASSIM can use any of the standard tools, such as minimizing the Akaike Information Criteria (AIC), Bayesian Information Criteria (BIC), cross-validation error, or by utilizing *χ*^2^-tests [21]. For the biological example, our inclusion of the siRNA perturbations made the theoretical considerations unable to meet the criteria for AIC, as the cost *V*(.) was a function of data points with and without the possibility of estimated standard deviation. We instead used a heuristic criterion to find a balance between model fit and sparsity, where the first edge removal that yielded a cost increase larger than 0.05 times the cost standard deviation of the parameter sets was chosen. The parameter value of 0.05 was chosen after studying the core system behavior, and deciding to correspond to the sparsest model with a retained fit of the data (Fig. S3, 38 parameters removed). To evaluate the fit between model and unperturbed time series data, we used χ^2^-tests [21]. In the χ^2^-tests, the degrees of freedom were set to equal the number of data points, and the rejection limit was set to α = 0.05.

### Bootstrap of optimization and model selection sensitivity

The minimization of Eq. (3) is a complex problem, and the risk of optimization failure is large, Moreover, optimization has a stochastic element, and the outcome varies. We therefore tested whether the impact of any optimization failure, and differences in outcome, had a greater impact than the information embedded in the data. We performed 24 corresponding bootstrap experiments, where individual time-series- and knock data points were assigned a homogenously distributed cost weight *τ ∊*[0,1], as seen in Eq. 4.

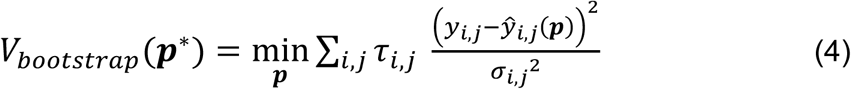

We found that the robustness of the parameter removal, in terms of removal order and final cost V_bootstrap_(***p***\*), was lower than the first comparison (Fig. S3). Even though approximately the same number of interactions could be removed, the chosen parameter set was different between different bootstrap experiments. This fact indicates that phenomena such as optimization failure have a nominal impact on model selection compared to the quality and certainty of our data.

### Time-series data from Th2 polarized total CD4+ T cells

Total CD4+ cells were isolated from PBMCs using isolation kits from Miltenyi (Bergisch Gladbach, Germany) according to the manufacturer′s instruction. The isolated cells were washed, activated and Th2 was polarized as previously described [35]. Briefly, the isolated total CD4+ cells were activated with plate-bound anti-CD3 (500ng/ml), 500ng/ml soluble anti-CD28, 5μg/ml anti-IL-12, 10ng/ml IL-4 and 17ng/ml IL-2 (R&D Systems). For microarray experiments, 200 ng total RNA was labeled and transcribed to cRNA using the Quick Amp Labeling Kit (Agilent, USA), then purified using the RNeasy mini kit and hybridized to Agilent Human GE 4x44 K v2 slides, according to the manufacturer′s instructions. Slides were scanned using the Agilent Microarray scanner 2505C and raw data were obtained using the Feature Extraction Software (Agilent). Moreover, we only considered data for genes that were deemed to respond to the data, so that we could reject the changes of expression over time from being white measurement, i.e. reject if the sum of residuals from the null-model > χ^2^_0.95_(df = 4).

### *In vitro* knock-down experiments

In vitro knock-downs were generated from human naïve CD4+ T-cells isolated from healthy donor buffy coats using magnetic bead separation (Miltenyi Biotec, Sweden). Typically, 1x10^6^ cells were transfected with 600nM on-target plus SMART pool against IRF4 (Dharmacon, USA), MAF, GATA3, USF2, COPEB (Thermo Fisher Scientific Inc, USA), or non-targeting siRNA (Dharmacon, USA) using Amaxa transfection (U-014). After six hours of incubation, the cells were washed and subsequently activated with plate-bound anti-CD3 (500 ng/ml), soluble anti-CD28 (500 ng/ml) and polarized towards Th2 using IL-4 (10 ng/ml), IL-2 (17 ng/ml) and anti-IL-12 (5 μg/ml, R&D Systems, USA) for 24 hours for each of the TFs siRNA (12h for IRF4). The cells were then harvested in Qiazol and total RNA was extracted using the RNeasy mini kit (QIAGEN, Germany). Expression was analyzed using the Human GE 4x44K v2 Microarray Kit from Agilent Technologies. All siRNA-induced knock-downs were followed by a short Th2 polarization (24h for all except IRF4, which had 12h) and microarray analysis, and control experiments corresponding Th2 polarization using scrambled non-targeting siRNA.

### *In silico* knock-downs

To overcome possible hurdles with dynamics outside the prediction ability of the model, and thus any bias in model fit, the data was truncated between −2 and 2. The knock-downs can be *in silico* modeled in a great variety of ways, and in LASSIM we chose to model homozygotic and heterozygotic knock-downs by increasing the degradation constant in Eq. (2), such that:

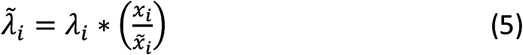

In Eq. 5, the augmented *λ* is estimated from the inverse of the data fold change. For each of the *in silico* siRNA knock-downs, 12 independent simulations were performed. The perturbations were approximated by increasing the self-regulatory term λ for each of the studied TFs. This increase was set to be proportionate to the measured decrease of said knocked gene.

## Data access

All utilized data are deposited at gene expression omnibus under the supe-series GSE60683, which will be publicly release upon acceptance. Reviewers can download the data using the private link https://www.ncbi.nlm.nih.gov/geo/query/acc.cgi?token=gxmneaiehlqxlar&acc=GSE60683.

## Supplementary material

**Fig S1. The time scale of LASSIM.** It was found that the time needed for the search algorithm to find a parameter solution that passes a χ^2^-test increased exponentially with number of parameters.

**Fig S2: LASSIM improved dynamic trajectories and structure compared to LASSO on two computer-generated examples.** (A) Simulations using LASSIM (dashed blue line) and LASSO (red solid lines) derived models on a synthetic dataset. The black dots and lines shows sampled and real expression of IRSP over time. LASSIM is able to identify the overshoot of IRSP between the first and second time-point correctly, whereas LASSO cannot. (B) LASSIM (red dots) receives higher AUC than our previous best performer LASSO method (blue line) in the DREAM2 network identification challenge in terms of precision [51]. LASSIM backward selection using the linear transfer function was applied to all connections with confidence score of >80% from our previous approach, and therefore has no predictions for recall>0.22. The area under the curve (AUC) of the receiver operating curve (ROC) was about 20% higher for LASSIM than for LASSO (P<0.03).

**Fig S3: Sensitivity analysis of core model identification.** When removing edges from the model, the randomness imbedded in the stochastic optimization will give rise to a difference in the system behavior (blue lines). To test if such optimization artifacts have a larger impact on the model selection than the quality of the data, a sensitivity analysis was performed, where each data point was given a uniformly distributed random weight between 0 and 1. The weighted version of the goodness-of-fit as a function of removed edges are plotted in red. As seen, the changes in the data weights induce a higher uncertainty to the core model identification. Moreover, the unperturbed model starts to rise around 37 removed parameters, which was used when determining the mu-coefficient in Eq. 4.

**Fig S4: The impact of knock data on the core model inference.** We tested the benefit of adding the siRNA knock down data as perturbations in the model inference, and found that the stability of the network inference dramatically increased. On the left side graphs, the model behavior without the perturbation data is shown. The results can be compared with the more stable version on the right, where knock data have been used.

**Fig S5: The fit of the random core models to the naïve data** The models that were found to fit the total t-cell data better than our prediction, were tested at the original training data, i.e. the naïve t-cell data. It was found that none could fit the data better than the prediction. Nevertheless, none of the models could be rejected using a χ^2^-test.

**Fig S6: Distribution of the certainty of the peripheral genes** The distribution of the fits of the peripheral genes on the total T-cell expression time series are shown. Here, the x-axis shows the goodness of fit (as calculated by Eq. 3), and the y-axis correspond to the frequencies.

**Fig S7: The saturation effect of the input to each gene** The nodes were modeled as nonlinear dependent on each other. Shown is the potential values of the input from other nodes. More specifically, the values are plotted such that they cover the range of the parameter search span for the minimal case of only one input to a transcription factor.

